# Regulating Cell Orientation with a Femtosecond Laser-Induced Macro Stripe Design on Metallic Culture Surfaces

**DOI:** 10.1101/2023.08.30.555452

**Authors:** Chikahiro Imashiro, Atsushi Ezura, Takahiro G. Yamada, Yoshikatsu Akiyama, Jun Komotori

## Abstract

Controlling cell orientation is paramount in bioengineering processes. While several surface modification techniques have emerged to guide cell alignment, they often involve complex, repetitive procedures for each experiment. A streamlined approach for cell orientation is thus necessary. In this study, we present a reusable metallic culture surface that induces an anisotropic cell orientation, attributed to its unique geometric morphology. By employing a femtosecond laser, periodic nanostructures were produced on the metallic culture surface, leading to a distinctive macro-stripe design composed of both laser-treated and mirrored areas. Myoblast cells cultured on this surface displayed a pronounced orientation. Initial random cell adherence was followed by migration towards the mirror surface, culminating in the desired orientation. This shift can be credited to the mirrored sections offering superior cell adhesion and reduced wettability compared to the laser-treated sections. This innovative culture surface holds significant potential for advancing bioengineering endeavors, especially in the realm of tissue engineering.

## Introduction

The regulation of cell orientation is pivotal in bioengineering processes such as tissue engineering, regenerative medicine, drug screening, and cultured food production. Anisotropic tissues play crucial roles [1]. For instance, in muscle tissue, the differentiation rate and cell tension can be enhanced by modulating cell orientation [2]. While numerous studies have explored cell orientation regulation using various techniques, many have focused on surface modification technologies [3]. These studies often prepared surfaces with varying wettabilities in stripe patterns on the culture medium, observing that cells typically favored hydrophobic surfaces, aligning according to the stripe orientation. Despite the availability of effective protocols to guide cell orientation, each experiment necessitates intricate workflows to engineer the culture surface, underscoring the need for a more streamlined method.

Current strategies largely hinge on the surface modification of disposable culture vessels, leading to labor-intensive workflows [4]. If a smart culture surface, capable of directing cell orientation, could be reused, each experiment would not entail such exhaustive preparation. Additionally, avoiding disposable vessels and repeated surface treatments would offer both cost and environmental benefits. As such, there’s a pressing need to refine and advance the bioengineering workflows for both researchers and industries. In recent work, we introduced reusable culture vessels crafted from biocompatible materials like titanium and SUS316L, commonly used in the human body [5–7]. These materials not only show excellent cytocompatibility but also demonstrate robust heat resistance in autoclaves, a prevalent sterilization technique. Nonetheless, the conventional approach to manipulate wettability involves surface treatments with chemical or biochemical agents [8]. These agents undergo changes during repeated use, especially in autoclaves, negating the benefits of reusable metallic culture surfaces.

Earlier studies have tailored the wettability of metallic culture surfaces using surface morphology, maintaining stability even in an autoclave [7,9–11]. Building on this, we present a novel approach: employing stripe-patterned metallic culture surfaces with differential wettabilities, derived from varied surface morphologies, to guide cell orientation. In our research, we textured biomedical-grade Ti–6Al–4V alloy surfaces to create a laser-induced periodic surface nanostructure (LIPSS) using femtosecond laser treatments, optimizing wettability [12]. Here, we developed reusable smart culture surfaces that regulate cell orientation, combining the LIPSS structures with polished mirror finish. To validate the efficacy of our approach, we used mouse myoblasts, a representative cell line for studying cell orientation.

## Results and discussions

### Design and evaluation of the culture surface

Drawing insights from prior research involving mammalian muscle cells [1], we adopted a stripe pattern combining 40 μm of a polished mirror surface with 40 μm of the LIPSS surface, as depicted in Fig. 1A. We labeled these culture surfaces the “PL-series”, integrating both polished mirror and laser-treated stripes. To evaluate the impact of this striped design, we fashioned two sample types: one exclusively with a polished mirror surface (referred to as the P-series) and the other with a solely laser-treated surface (L-series). Comprehensive illustrations and evaluations of these samples are provided in Figs. 1B-E. Figures 1B-D present the field-emission scanning electron microscopy (FE-SEM; JSM-IT700HR, JEOL) images of the PL-, P-, and L-series respectively. The PL-series stripes’ width was measured to be 40.6 ± 1.7 μm, with the laser-treated surfaces presented in 78 μm periodic cycles. This slight discrepancy in measurement stemmed from variations in the working distance between the laser equipment and the sample pieces. Fig. 1C showcases the pristine mirror surface typical of the P-series, while Fig. 1D depicts a nanogroove structure on the L-series’ surface. This nanogroove structure exhibited a periodicity of approximately 300 nm.

**Figure 1.**
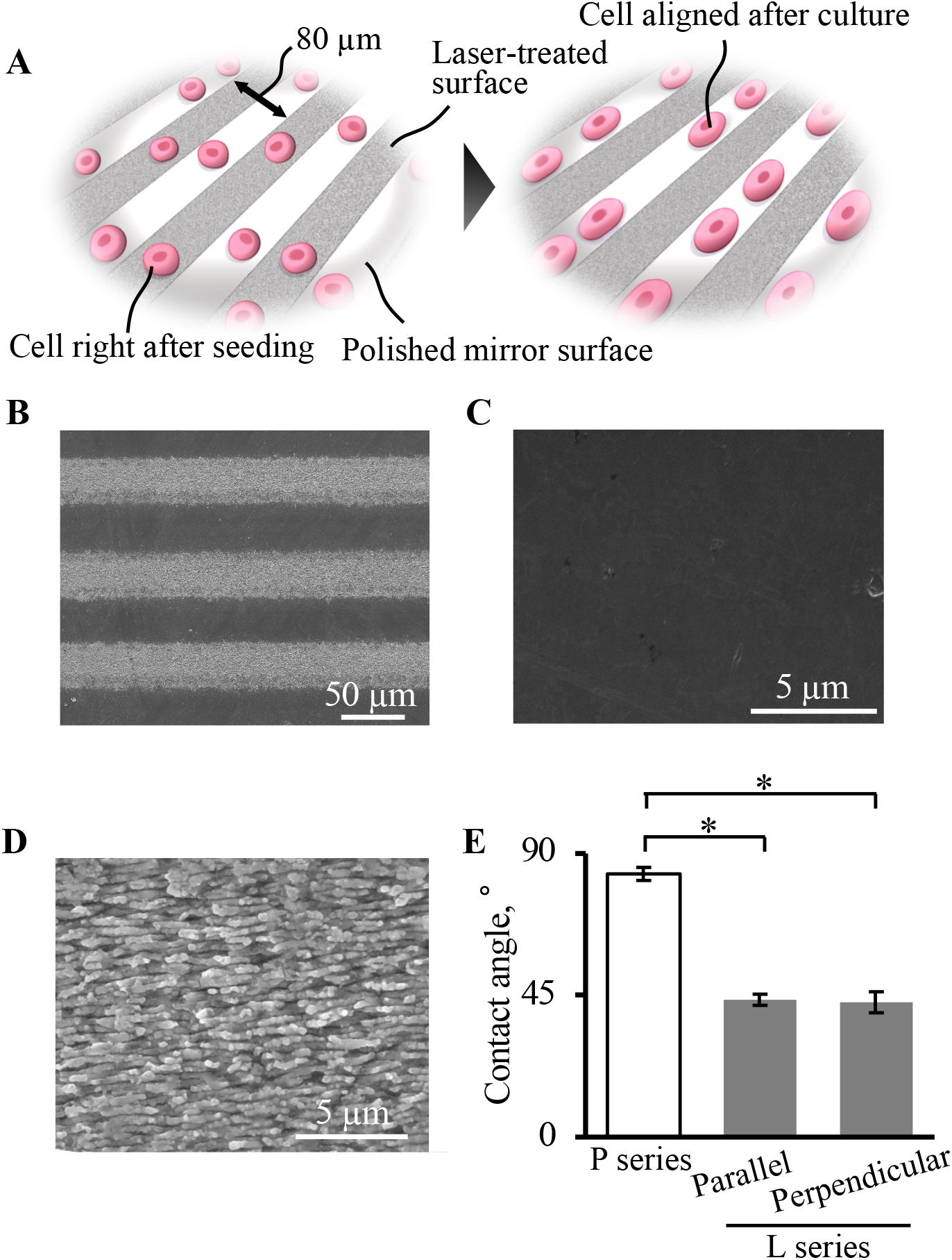
Design and Assessment of the Culture Surface. A: Illustration and rationale for the culture surface. The mutual stripe design consists of 40 μm wide laser-treated surfaces adjacent to 40 μm mirror surfaces. This design encourages cells to migrate toward the mirror surface, achieving alignment. C-E: SEM images showcase the morphological details of the PL-, P-, and L-series surfaces. E: Contact angle measurements for the P and L-series. For the L-series, angles were measured both parallel and perpendicular to the laser scanning direction. Statistical significance was tested using Welch’s t-test (*N*=5, *adjusted *p*-value<.05, represented as mean±SD).

Furthermore, Fig. 1D presents the contact angle measurements for both the P and L-series. For the L-series, readings were taken at two orientations: parallel and perpendicular to the laser-scanning direction. Notably, laser processing considerably augmented the surface’s wettability, signaling diminished cell adhesion to the LIPSS surface. This wettability enhancement can be attributed to morphological and chemical modifications of the culture surface [13].

### Cell orientation and activities on the culture surface

Figure 2 depicts the orientation (A-B) and other activities (C-D) of cells cultivated in the different series. Cell orientation was evaluated with cells seeded at a density of 5.0 x 10^3^ cells/cm^2^. Fluorescence images of the cells, stained using calcein-AM (C396, Dojindo), are displayed in Fig. 2A. The images reveal that the PL-series effectively directed cell orientation. This observation was quantified utilizing the ImageJ software (version 2.10/1.53c), with the resultant angle distribution graphically represented in a rose diagram, as seen in Fig. 2B. According to this data, the PL-series consistently aligned cells in accordance with the stripe design across all culture durations (detailed statistical analysis can be found in Table 1). Conversely, the L-series began demonstrating notable cell orientation after 72 h of cultivation. This aligns with existing literature, suggesting the LIPSS surface’s capacity to guide cell orientation [14]. These findings underscore that a microscale stripe design integrated with a nanoscale structure is more adept at cell alignment compared to a continuous nanostructure.

**Table 1.** Circular Statistics and Rayleigh Test Results for Every Experimental Condition.

| Condition | Duration | Circular Mean* | Kappa | p.value | adjusted p. value** | Significance (adjusted p.value < 0.05) |
| --- | --- | --- | --- | --- | --- | --- |
| P series | 24 h | 1.40252715 | 0.14340536 | 6.28E-02 | 5.66E-01 | NO |
|  | 48 h | 1.37891709 | 0.10367068 | 8.94E-02 | 8.05E-01 | NO |
|  | 72 h | -0.94383556 | 0.03518778 | 4.70E-01 | 1.00E+00 | NO |
| L series | 24 h | -1.28583012 | 0.13174838 | 6.98E-02 | 6.29E-01 | NO |
|  | 48 h | -1.47293125 | 0.07519623 | 2.19E-01 | 1.00E+00 | NO |
|  | 72 h | 0.19797142 | 0.38167569 | 3.56E-19 | 3.20E-18 | YES |
| PL series | 24 h | -0.09961872 | 0.90920256 | 3.06E-29 | 2.75E-28 | YES |
|  | 48 h | -0.23659269 | 0.67449738 | 8.63E-22 | 7.76E-21 | YES |
|  | 72 h | -0.08663317 | 0.53865509 | 2.67E-69 | 2.41E-68 | YES |

\* Angular units are expressed in radians
\*\* p.value after FWER control with Bonferroni correction

**Figure 2.**
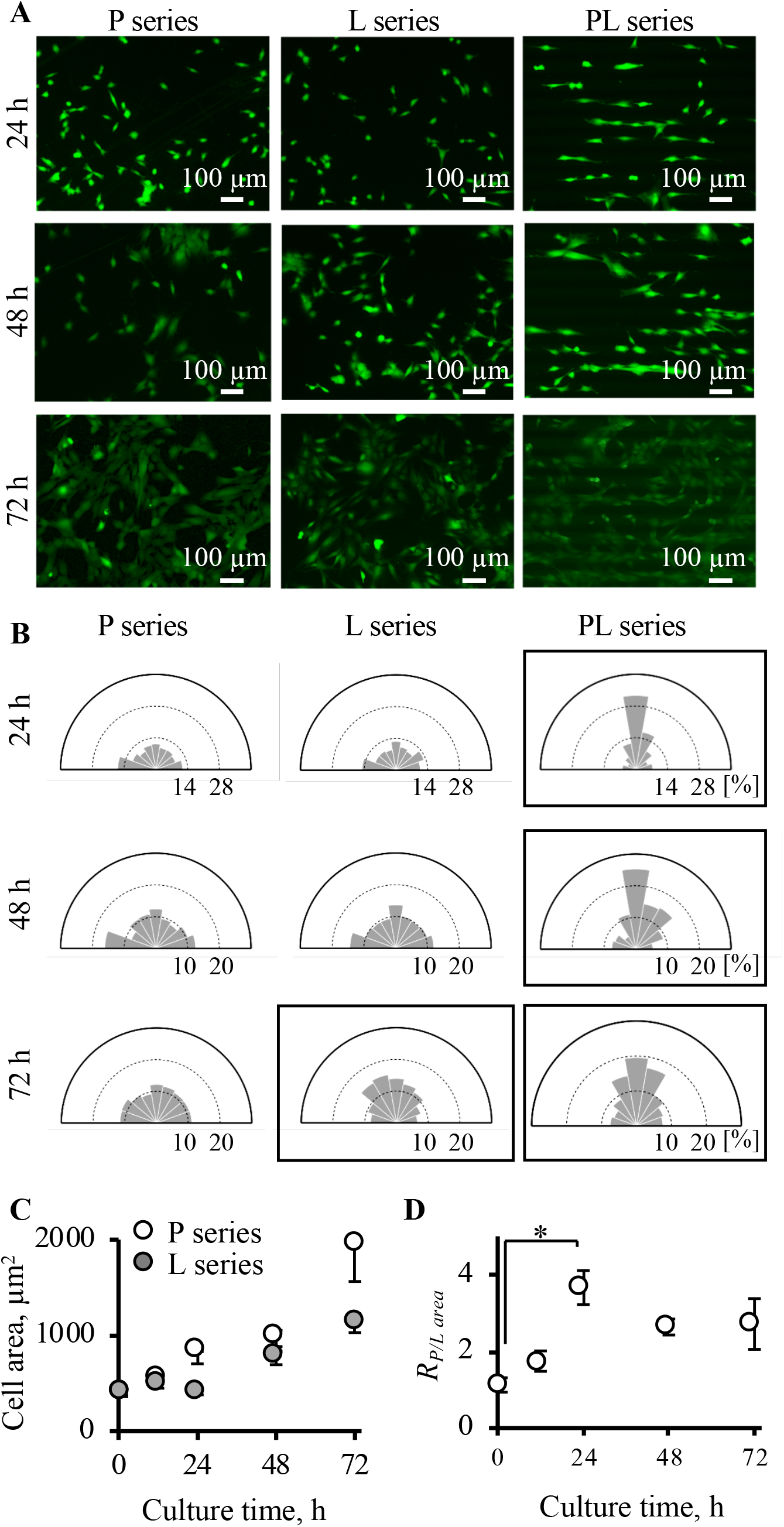
Evaluation of Cell Orientation and Activity on Different Culture Surfaces. A: The orientation of live cells, stained with Calcein, is depicted. B: The directional distribution of cells is illustrated in a rose diagram, segmented into 9 bins, each spanning 20°. Graphs showing statistical significance are highlighted with squares (adjusted *p*-value<0.05). Experiments under each condition were conducted thrice, varying both the device and cell samples, and their results are consolidated in this figure. Refer to Supplementary Table 1 for the count of evaluated cells in every sample. C: Quantification of the area occupied by each cell on the P and L-series. Refer to Supplementary Table 2 for the count of evaluated cells in every sample. D: For the PL-series, the distribution of cells is evaluated. The *R*_*P/L area*_ metric, which denotes the ratio of cell coverage on the polished mirror surfaces to the LIPSS surfaces, provides insights into the temporal evolution of cell distributions. Experiments for each scenario were conducted three times, utilizing different devices and cell samples. Their results are summarized in this figure. The significance was assessed using Welch’s t-test (*N*=3, *adjusted *p*-value<.05, depicted as mean±SD).

While the primary focus of this research was on the utilization of the tailored surface as a culture vessel, the design holds potential for broader bioengineering applications. Biomaterials, like artificial joints requiring cell alignment, could benefit from these engineered surfaces [14,15]. Moreover, the insights gained from our research can enrich the development of lab-on-a-chip devices employing aligned cells [16,17]. An additional advantage is that the PL-series demands less laser processing time and effort compared to the L-series. It’s also worth noting that, since this study capitalizes on geometric cues from mechanical modifications, integrating this innovative and sustainable culture surface with other methodologies employing ultrasound, polymer, protein coatings, or shear stress becomes feasible [18–23].

Our working hypothesis proposes that, due to varying adhesion properties of culture surfaces, cells exhibit a preference for polished surfaces which possess greater adhesiveness, leading to the alignment of cells. To validate this hypothesis, we examined the cellular response to both laser-treated and mirrored surfaces utilizing fluorescent imagery, as depicted in Figs. 2C-D. Cells were seeded at a density of 1.0 x 10^3^ cells/cm^2^. Figure 2C presents a quantitative analysis of cellular areas in both the P- and L-series, accomplished through image processing in ImageJ, focusing on calcein-stained cells. As evident in Fig. 2C, the P-series exhibited a more pronounced increase in cellular area when compared to the L-series. Moreover, variations in both series and culture duration significantly influenced alterations in cellular areas (refer to Supplementary Table 3). Given that cell adhesion force directly correlates with cell area [24], the P-series manifested superior cell adhesion. Consequently, our findings not only inferred from wettability but also from direct evaluation of cell adhesiveness, substantiating the reduced cell adhesiveness associated with the LIPSS on the Ti–6Al–4V surface in this study.

Additionally, both series did not reveal disparities in cellular areas during the initial phase of culture. This suggests that variations in cell adhesion forces might emerge after a specific culture period. We theorized that cells initially adhere at random to the PL-series’ culture surface, subsequently developing a predilection for mirror over LIPSS surfaces over a certain duration. As a result, cells begin migrating from laser-treated regions to mirrored ones. This theory was further scrutinized by examining cell distribution within the PL-series, as illustrated in Fig. 2D. For each time checkpoint, cells underwent calcein-AM staining, and the cell coverage area ratio was determined. In Fig. 2D, the y-axis denotes the *R*_*P/L area*_, or the ratio of the cell-covered area on polished mirror surfaces compared to that on LIPSS surfaces. The graph indicates an initial random cellular adhesion to the PL-series, with a subsequent shift towards the polished mirror surface. Intriguingly, the *R*_*P/L area*_ peaks at 24 hours, with the sole significant *R*_*P/L area*_ discrepancy at 1 hour being evident at this 24-hour mark. This suggests that, following local confluence on the PL-series’ polished area, cells expanded into the LIPSS region. Such data implies the potential for cultivating a uniformly aligned cell monolayer either through prolonged culture or amplified seeding. If successfully generated and harvested, this aligned cell monolayer could significantly benefit cell sheet technology [25]. Various cell harvesting techniques have been explored, some of which are compatible with metallic culture surfaces [26,27]. Additionally, this principle is not confined to linear stripes but can be extrapolated to curves or intricate patterns, offering considerable potential for advancements in tissue engineering.

## Conclusion

We introduced a microscale stripe design combined with nanoscale grooves on metallic culture surfaces. Our findings illustrate that this integrated design more effectively directs cell orientation than solely utilizing nanoscale grooves. Upon exposure to the newly developed culture surface, cells initially adhered in a random fashion, later migrating towards the mirror surfaces, resulting in a distinct cell alignment. Moreover, once local confluence was achieved on the mirror surface, cell growth expanded to the LIPSS surface, suggesting the potential to form a complete cell layer. This innovative design paves the way for advanced applications in tissue engineering. Additionally, the metallic culture surface offers the advantage of being reusable via autoclave sterilization, and its geometric morphology remains intact, benefiting from the inherent durability of metallic composition.

## Materials and methods

### Culture surface fabrication

We utilized the Ti-6Al-4V ELI alloy (ASTM F136) for our sample material. Each sample measured 15 mm in diameter and 3 mm in thickness. Before undergoing laser treatment, the surfaces of these samples were meticulously polished using emery paper and then refined with colloidal silica slurry to achieve a mirror-like finish. The prepared samples were exposed to an ytterbium-fiber femtosecond pulsed laser (FCPA μ Jewel 02, I-D-6-03, IMRA AMERICA, INC., Michigan, USA). The laser, having a wavelength of 1041.5 nm, operated with a pulse duration of 367 fs, a frequency of 200.9 kHz, a fluence of 0.401 J/cm^2^, a scan rate of 9 mm/s, and was utilized with two distinct hatching pitches. For the L-series fabrication, a hatching pitch of 40 μm was employed during laser exposure. This process led to the formation of LIPSS across the entire sample surface. In contrast, the LP-series was crafted using a laser hatching pitch of 80 μm, resulting in a distinct stripe pattern. Subsequent to the laser treatment, the surfaces were closely examined with FE-SEM to elucidate the intricacies of their micro- and nanometer-scale morphologies.

### Contact angle measurement

We employed the sessile drop technique to measure static contact angles. Throughout the measurement process, humidity was consistently maintained between 40% and 60%. The contact angle of a water droplet, placed perpendicular to the culture surface, was ascertained using the θ/2 method. This analysis was conducted with the aid of a contact angle meter (DSA100, DAS3 software, KRÜSS GmbH, Hamburg, Germany) [28].

### Cell culture

For all experiments, we utilized mouse myoblasts (C2C12) sourced from the Riken Bio Resource Center (RCB0987; Ibaraki, Japan). These cells were cultivated in a growth medium inside a 5% CO_2_ humidified incubator maintained at 37°C. When passaging was necessary, cells underwent trypsinization using a 0.05% trypsin-EDTA solution (Product No. 25300; Life Technologies, Carlsbad, CA, USA).

### Statistical analysis

We utilized Welch’s t-test to discern differences between any two groups. For evaluations involving multiple groups, the p-value was corrected using the Bonferroni method to identify significant disparities, with a significance threshold set at *α*=0.05.

To ascertain the presence of cell orientation, we employed the Rayleigh test for uniform distribution based on the Von Mises distribution [29]. All derived p-values for each test condition underwent Bonferroni correction. We confirmed adjusted *p*-values against a significance level of *α*=0.05. For testing purposes, we leveraged the *r*.*test* function from the CircStats package [30] in R. The ggplot2 package [31] in R facilitated the creation of rose diagrams.

To probe the combined influence of culture time and the variance between the P and L-series on alterations in cell adhesion area, we devised a generalized linear model. This model encompassed interaction effects and presupposed a gamma distribution. The significance of the factors’ effects was ascertained via a two-way analysis of variance (ANOVA), implementing a likelihood ratio test for each factor at the *α*=0.05 significance level. These computations were executed using the *glm* and *anova* standard functions in R.

## Acknowledgment

The authors thank Mr. Kei Takahashi, Mr. Motohiko Takeuchi, and Mr. Junya Matsuzaki at Keio University for their support for experiments. This work was supported by JSPS KAKENHI Grant Number 22K18188 and grants from the Light Metal Educational Foundation, Inc., the Japan institute of Metals and Materials, and Nissin Sugar Co., Ltd.

