## Supplemental Information for "Regulating Cell Orientation with a Femtosecond Laser-Induced Macro Stripe Design on Metallic Culture Surfaces"

Macro stripe design of femtosecond laser-induced periodical surface nanostructure and mirror surface on metallic culture surface strongly regulates cell orientation

Chikahiro Imashiro[a,b*], Atsushi Ezura[c], Takahiro G. Yamada[d], Yoshikatsu Akiyama[b], Jun Komotori[e**]

[a] Graduate school of Engineering, The University of Tokyo, Tokyo, Japan

[b] Institute of Advanced Biomedical Engineering and Science, Tokyo Women's Medical University, Tokyo, Japan

[c] Faculty of Engineering, Sanjo City University, Niigata, Japan

[d] Department of Molecular Biology, University of California San Diego, La Jolla, CA, USA.

[e] Department of Mechanical Engineering, Faculty of Science and Technology, Keio University, Kanagawa, Japan

Supplementary Table 1. The number of evaluated cells in each sample for Fig. 2B.

| Condition | Duration (h) | Replicate No. | Number of Cells |
|---|---|---|---|
| L series | 24 | 1 | 283 |
| | | 2 | 167 |
| | | 3 | 166 |
| | 48 | 1 | 598 |
| | | 2 | 157 |
| | | 3 | 321 |
| | 72 | 1 | 145 |
| | | 2 | 532 |
| | | 3 | 532 |
| P series | 24 | 1 | 101 |
| | | 2 | 221 |
| | | 3 | 219 |
| | 48 | 1 | 362 |
| | | 2 | 387 |
| | | 3 | 152 |
| | 72 | 1 | 448 |
| | | 2 | 268 |
| | | 3 | 1,725 |
| PL series | 24 | 1 | 99 |
| | | 2 | 140 |
| | | 3 | 144 |
| | 48 | 1 | 172 |
| | | 2 | 96 |
| | | 3 | 207 |
| | 72 | 1 | 332 |
| | | 2 | 1,029 |
| | | 3 | 974 |

Supplementary Table 2 The number of evaluated cells in each sample for Fig. 2C.

| Condition | Duration (h) | Replicate No. | Number of Cells |
|---|---|---|---|
| L series | 1 | 1 | 23 |
| | | 2 | 46 |
| | | 3 | 43 |
| | 12 | 1 | 44 |
| | | 2 | 27 |
| | | 3 | 40 |
| | 24 | 1 | 146 |
| | | 2 | 161 |
| | | 3 | 82 |
| | 48 | 1 | 92 |
| | | 2 | 167 |
| | | 3 | 166 |
| | 72 | 1 | 598 |
| | | 2 | 157 |
| | | 3 | 321 |
| P series | 1 | 1 | 61 |
| | | 2 | 63 |
| | | 3 | 50 |
| | 12 | 1 | 50 |
| | | 2 | 55 |
| | | 3 | 57 |
| | 24 | 1 | 146 |
| | | 2 | 161 |
| | | 3 | 82 |
| | 48 | 1 | 92 |
| | | 2 | 115 |
| | | 3 | 103 |
| | 72 | 1 | 107 |
| | | 2 | 77 |
| | | 3 | 92 |

Supplementary Table 3 The results of the statistical analysis in Fig. 2C

| Factor | Df | Deviance | Resid. Df | Resid. Dev | Pr(>Chi) | Significance |
|---|---|---|---|---|---|---|
| time | 1 | 631.22 | 2922 | 2325.2 | <2.2E-16 | YES |
| PL | 1 | 112.88 | 2921 | 2212.4 | 1.40E-06 | YES |
| time:PL | 1 | 11.54 | 2920 | 2200.8 | 0.1229 | NO |
